# Phytoformic Gold in Ash Samples of Plants from the North Goa Iron Ore Mining Belt: Detection, Characterisation, X-ray Diffraction, and Spectroscopic Evidence for Biogeochemical Gold Nanoparticle Formation

**DOI:** 10.64898/2026.05.15.725495

**Authors:** Naik Shruti, N.M. Kamat

**Affiliations:** Botany program, Mycolab, SBSBT, Goa University, Taleigao, Goa – 403 206, India

**Keywords:** phytoformic gold, phytoauroliths, gold nanoparticles, Debye–Scherrer, XRD, FTIR, chloroauric acid, Banded Iron Formation Goa, Western Dharwad Craton, phytoextraction, biogeochemical prospecting

## Abstract

Gold is widely distributed in the biosphere, and higher plants growing on geochemically anomalous substrates can accumulate significant amounts of gold. This study reports, for the first time from Goa, the detection, spectroscopic characterisation, and X-ray diffraction analysis of phytoformic gold — biologically sequestered crystalline gold — in the above-ground dry litter ash of six tree species *(Acacia auriculiformis, Alstonia scholaris, Anacardium occidentale, Artocarpus heterophyllus, Ficus benghalensis, Syzygium cumini*) growing on mining dumps within the North Goa Banded Iron Formation (BIF) Belt of the Western Dharwad Craton. Microgravimetric analysis of aqua regia-extracted heavy ash fractions revealed gold concentrations of 275–1100 ppm, two to five orders of magnitude above the crustal background (∼0.004 ppm). Fourier Transform Infrared (FTIR) spectroscopy of 0.22□μm membrane-filtered crude extracts confirmed the tetrachloroaurate(III) complex [AuCl□]□ as the dominant dissolved gold species, with the diagnostic 1400–1700□cm□^1^ absorption envelope present in all six species. UV–Visible spectrophotometry confirmed chloroauric acid formation with a universal λmax at 372.5□nm across all species. Powder X-ray diffraction (XRD) of heavy ash fractions yielded the characteristic FCC metallic gold reflections Au(111), Au(200), and Au(220) in all five species analysed. Application of the Debye–Scherrer equation to the Au(111) reflection (2θ = 38.2°, Cu Kα) established crystallite sizes of 17.7–31.8□nm, confirming that phytoformic gold exists as nanoscale crystalline particles in all species. *Ficus benghalensis* produced the largest and most crystalline gold nanoparticles (31.8□nm) and uniquely exhibited strawberry-shaped isomorphic auriferous siliceous biominerals designated ‘phytoauroliths’. The described low-cost protocol — ashing, aqua regia extraction, membrane filtration, and multi-technique spectroscopic and diffraction confirmation — constitutes a validated method for rapid biogeochemical gold anomaly detection. Applications in gold phytoextraction and mining waste phytoremediation are discussed.

## 1. Introduction

Gold has been detected in plant tissues since the early twentieth century. In 1900, the fire-ashing method was first applied to hardwood trees, yielding visible gold beads and establishing the biological concentration of gold as a quantifiable phenomenon (Girling and Peterson, 1980). Subsequent investigations have demonstrated that gold can be solubilized from minerals and soils through microbial activity and cyanogenic metabolites produced by roots and rhizosphere microorganisms. Once taken up by plants, they become organically bound and transported to protective and structural tissues, ultimately accumulating in organs that are periodically discarded, including leaves and rootlets (Jones et al., 1989; Christopher et al., 1999).

Gold, which is taken up, transported, biologically reduced or transformed, and deposited as a morphologically distinct biomineral exhibiting diverse associations, shapes, and sizes, is defined in the present work as *phytoformic gold*. This term encompasses nano-to microsized gold particles that retain biological imprints of their formation environment, distinguishing them from abiogenic or thermochemically processed gold particles.

The Western Dharwad Craton of peninsular India, exposed across the iron ore mining districts of North Goa, constitutes a classic Archaean greenstone terrane characterized by Banded Iron Formation (BIF) / Banded Hematite Quartzite (BHQ) lithologies, auriferous quartz veins, and gold– sulfide mineralization. Mining dumps and revegetated waste embankments within this geological province represent unusual pedological settings in which plant species intercept geochemically anomalous gold concentrations in the rhizosphere of plants.

Despite the extensive global literature on phytoextraction and biogeochemical prospecting for gold, no systematic study documenting phytoformic gold in the vegetation of Goa has been published prior to the present investigation. This study reports the first confirmed detection and multi-technique characterization of phytoformic gold in the above-ground litter of six tree species from North Goa mining dumps. Crucially, for the first time in this system, powder XRD and Debye– Scherrer crystallite size analysis confirmed that gold in these plant tissues existed as crystalline nanoparticles, establishing their classification as biogenic gold nanoparticles (bio-AuNPs) of phytogenic origin.

## 2. Materials and Methods

### 2.1 Study Area and Plant Species

The study area is located within the North Goa Banded Iron Formation Belt of the Western Dharwad Craton. Six tree species abundant on active and abandoned mining dumps were selected: *Acacia auriculiformis* A. Cunn. ex Benth. (Fabaceae), *Alstonia scholaris* (L.) R.Br. (Apocynaceae), *Anacardium occidentale* L. (Anacardiaceae), *Artocarpus heterophyllus* Lam. (Moraceae), *Ficus benghalensis* L. (Moraceae), and *Syzygium cumini* (L.) Skeels (Myrtaceae).

### 2.2 Sample Collection and Ash Preparation

Well-dried aboveground leaf litter was collected from representative individuals of each target species on the mining dumps. The litter was dried to a constant mass at 70°C. Aliquots of 10□g (dry weight) were ignited in a muffle furnace (550–600°C, 4□h) to produce inorganic ash. The ash content was calculated as a percentage of the initial dry litter mass. The heavy fraction was recovered using density fractionation.

### 2.3 Aqua Regia Extraction

The heavy fraction of each ash sample was extracted with freshly prepared aqua regia (3 parts concentrated HCl : 1 part concentrated HNO□, v/v) under mild warming (60°C, 30□min) with constant stirring. After cooling, the extract was filtered through a 0.22□μm cellulose acetate membrane. The filtrate, a crude chloroauric acid solution, was retained for spectroscopic analysis. Slides of ash fractions with and without thermal treatment at 200°C were prepared for optical microscopy.

### 2.4 Microgravimetric Estimation of Gold

Crude gold content in the heavy fraction was estimated by microgravimetry following selective precipitation and expressed as ppm (μg□g□^1^) in the heavy ash fraction.

### 2.5 UV–Visible Spectrophotometry

Aqua regia extracts were scanned at 190–750□nm using a double-beam UV–Visible spectrophotometer. The spectra were compared with pre-recorded reference spectra of pure H[AuCl□]·4H□O and H[AuCl□] admixed with humic acid, and matching λmax values were identified.

### 2.6 FTIR Spectroscopy

Sub-samples of heavy ash fraction were pressed as KBr discs and scanned at 400–4000□cm□^1^ (resolution 4□cm□^1^). The 0.22□μm filtered aqua regia extracts were also analysed as liquid films. The spectra were processed using OriginPro v5.0. Absorption bands were compared with the reference spectrum of pure H[AuCl□] with and without humic acid, and the matching frequencies were tabulated.

### 2.7 Powder X-ray Diffraction (XRD)

Powder XRD patterns of the heavy ash fractions were recorded over 2θ = 20°–80° using a powder X-ray diffractometer with Cu Kα radiation (λ = 1.5406□Å). The observed reflections were matched against JCPDS reference cards for metallic gold (Au□, FCC, JCPDS 04-0784), calcite (JCPDS 05-0586), hematite (JCPDS 33-0664), magnetite (JCPDS 19-0629), and quartz (JCPDS 46-1045). The crystallite size of the phytoformic Au(111) phase was determined using the Debye–Scherrer equation:

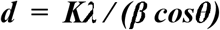

where *d* is the mean crystallite size (nm), λ =1.5406□Å (Cu Kα), *K*= 0.89 (Scherrer constant), β is the corrected full-width at half-maximum (FWHM) of the Au(111) reflection in radians after quadrature subtraction of instrument broadening (β_inst_ =0.10° in 2θ), and θ is the Bragg angle (half of the observed 2θ). An instrument-broadening correction was applied as 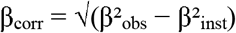.

### 2.8 Optical Microscopy

Ash fractions with and without thermal treatment at 200°C were mounted on slides and examined using optical microscopy. The morphology, size, and mineral associations of gold particles and biominerals were documented using photomicrography.

## 3. Results

### 3.1 Ash Content and Gold Concentration

The percentage ash content, heavy fraction proportion, and microgravimetrically estimated gold content for each species are listed in Table 1. The ash yields ranged from 29.77% (*F. benghalensis*) to 38.59% (*A. auriculiformis*). Gold content in the heavy fraction ranged from 275□ppm (*A. heterophyllus*) to 1100□ppm (*A. scholaris*), representing a biomagnification factor of ∼70,000– 275,000 relative to mean crustal gold (∼0.004□ppm).

**Table 1.** Ash content and microgravimetrically estimated gold content in the heavy fraction of dry litter biomass of six tree species, North Goa BIF Belt.

| Plant Species | Ash (%) | Heavy Fraction (%) | Gold in ash (ppm) | Rank by Au |
| --- | --- | --- | --- | --- |
| *Acacia auriculiformis* | 38.59 | 0.15 | 1050 | 2nd |
| *Alstonia scholaris* | 35.10 | 0.12 | 1100 | 1st |
| *Anacardium occidentale* | 31.10 | 0.04 | 675 | 4th |
| *Artocarpus heterophyllus* | 33.20 | 0.01 | 275 | 6th |
| *Ficus benghalensis* | 29.77 | 0.04 | 875 | 3rd |
| *Syzygium cumini* | 32.21 | 0.09 | 450 | 5th |

### 3.2 Morphology of Phytoformic Gold

Optical microscopy of the ash fractions revealed gold nano-and microparticles in all six species. Upon aqua regia treatment and mild heating, refined forms of gold particles and thin gold films were observed. The morphologies included isometric particles, irregular flakes, and dendritic forms. Strawberry-shaped isomorphic auriferous siliceous biominerals were consistently observed in the unprocessed ash of Ficus benghalensis. These are designated ***‘phytoauroliths’*** — a novel morphological category of phytoformic gold characterized by a biosilicon (auriferous quartz) association, which will be fully characterized in a forthcoming publication.

### 3.3 FTIR Spectroscopic Analysis

FTIR spectra (400–4000□cm□^1^) of the 0.22□μm filtered crude aqua regia extracts are presented in Figures 1–7. The dominant diagnostic feature of pure chloroauric acid in the mid-IR — a broad, intense absorption centred at 1400–1700□cm□^1^, attributable to the [AuCl□]□ complex ion, associated ionic species, and residual HNO□/NO□□ from the aqua regia medium — was present in all six plant species. A broad O–H envelope at 2800–3600□cm□^1^ was consistent with the strongly hydrated nature of H[AuCl□]·4H□O. The matching band data are presented in Table 2.

**Table 2.** Matching FTIR absorption bands (cm□^1^) between reference chloroauric acid + humic acid and crude aqua regia extracts of plant ash. +□=□band present; −□=□band absent.

| Ref. ($\text{cm}^{-1}$) | Acacia | Alstonia | Anacardium | Artocarpus | Ficus | Syzygium | Assignment (tentative) |
| --- | --- | --- | --- | --- | --- | --- | --- |
| 445 | + | – | + | + | – | – | Au–Cl def. |
| 540 | – | + | + | – | – | – | M–Cl str. |
| 675 | – | + | – | – | – | – | Fe–O/Au–Cl |
| 755 | + | + | + | + | + | + | M–Cl/M–O str. |
| 855 | + | + | + | + | + | + | M–Cl/M–O str. |
| 940 | – | – | – | – | + | – | Si–O–Si |
| 1040 | + | + | + | – | + | – | Si–O/P–O |
| 1130 | – | – | – | + | + | – | Si–O–Si asym. |
| 1340 | + | + | + | – | – | + | $\text{NO}_2$ (AqR) |
| 2340 | – | + | + | + | + | + | $\text{CO}_2/\text{NO}_2$ overtone |
| 2870 | + | – | – | + | + | – | C–H sym. str. |
| 2920 | – | – | – | – | – | – | C–H asym. str. |

**Figure 1.**
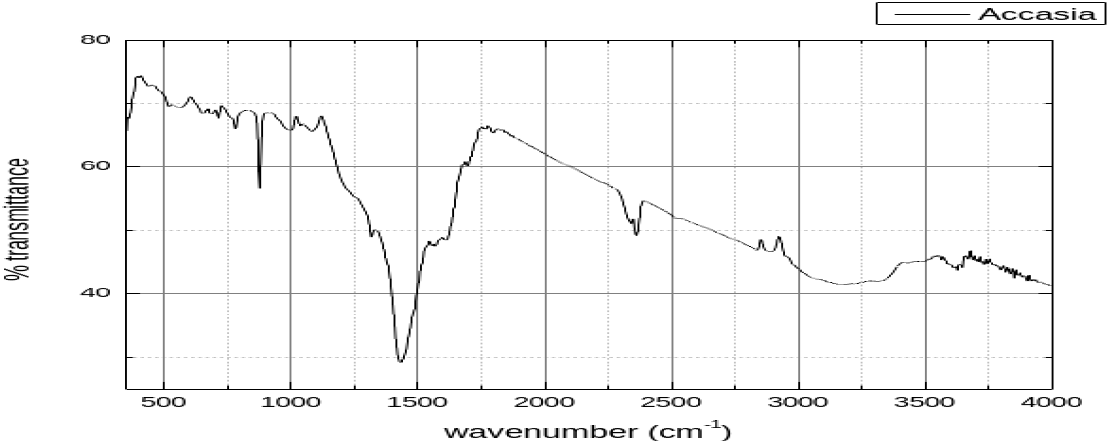
FTIR spectrum (400–4000 cm□^1^) of 0.22 μm filtered crude aqua regia extract of Acacia auriculiformis ash. The deep trough at 1400–1600 cm□^1^ and broad O–H envelope at 2800–3600 cm□^1^ are consistent with the presence of [AuCl□] □ ions.

**Figure 2.**
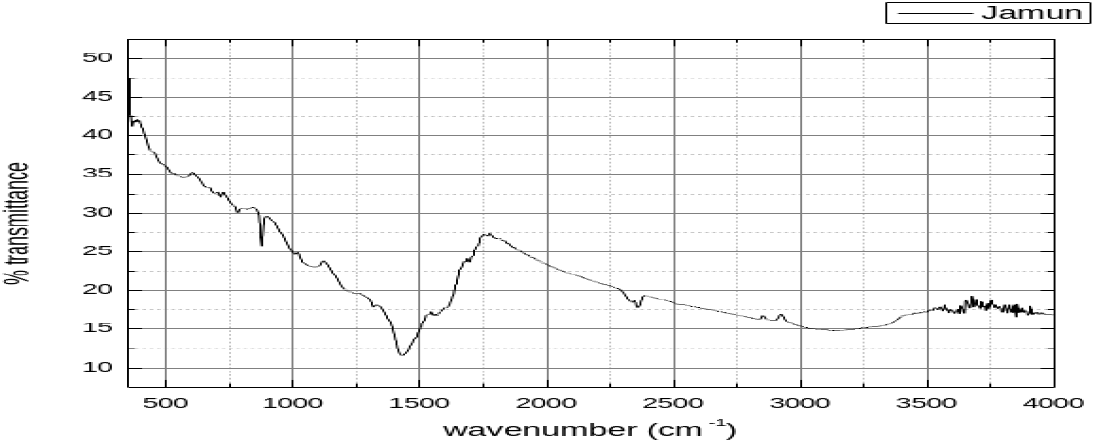
FTIR spectrum of crude aqua regia extract of Syzygium cumini (Jamun) ash. Low overall transmittance (10– 42%) reflects high dissolved solids, and the 1400–1600 cm□^1^ chloroaurate trough is well-resolved.

**Figure 3.**
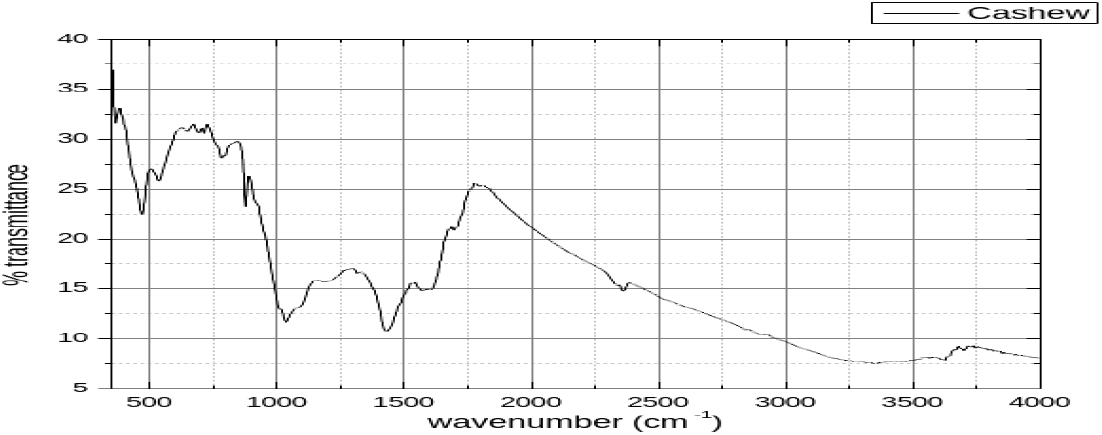
FTIR spectrum of crude aqua regia extract of Anacardium occidentale (Cashew) ash. The complex fingerprint region (400–1100 cm□^1^) reflects the silicate/phosphate-rich matrix, and the chloroaurate bands are partially masked.

**Figure 4.**
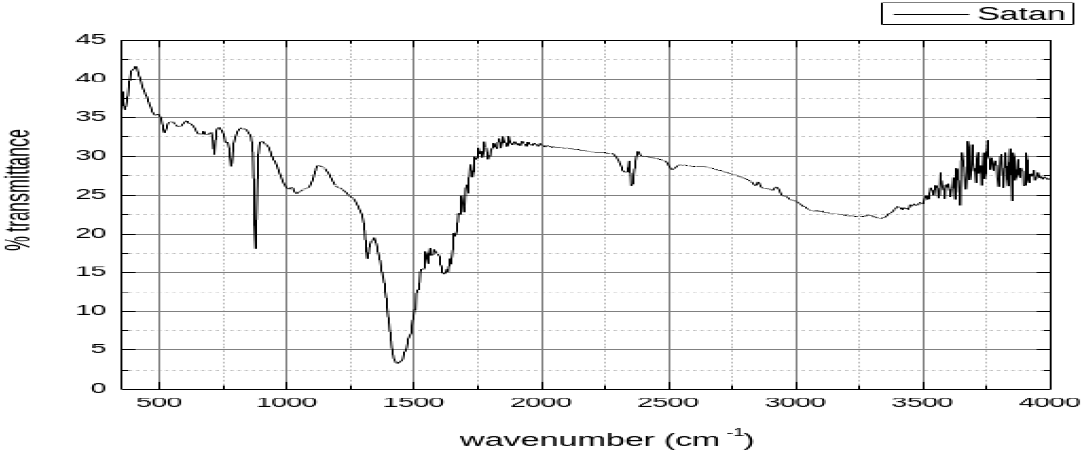
FTIR spectrum of crude aqua regia extract of Alstonia scholaris ash. The near-zero transmittance trough at ∼1400–1550 cm□^1^ is the strongest chloroaurate mid-IR absorption in the series.

**Figure 5.**
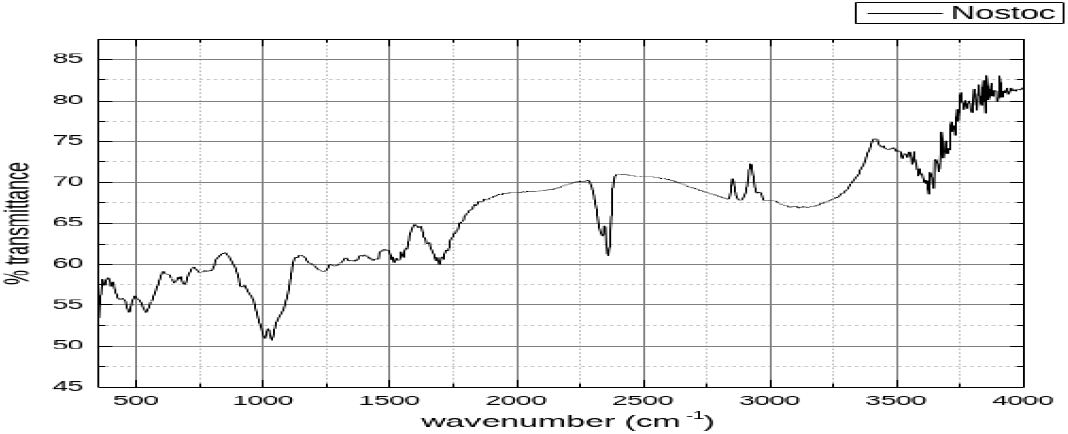
FTIR spectrum of Nostoc (cyanobacterial reference). The high overall transmittance (50–85%) and absence of the 1400–1700 cm□^1^ trough confirm the absence of chloroauric acid in this non-vascular phytobiont extract.

**Figure 6.**
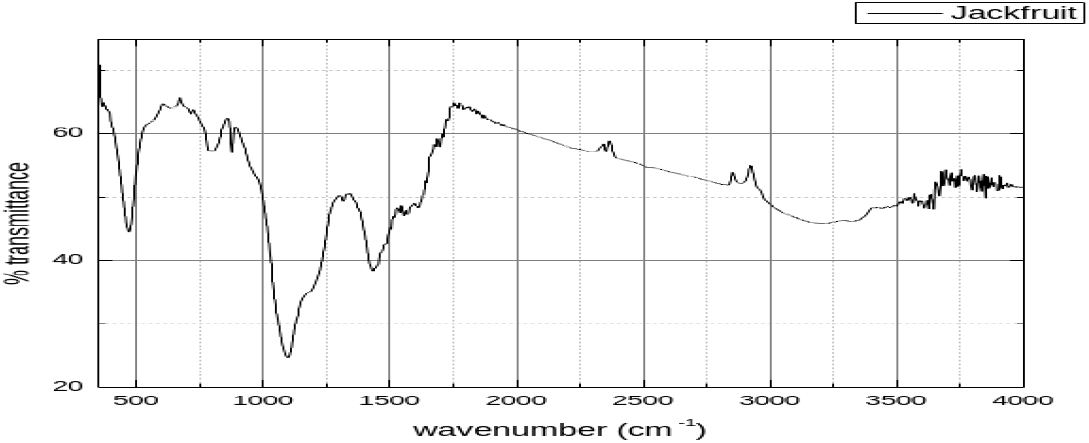
FTIR spectrum of crude aqua regia extract of Artocarpus heterophyllus (Jackfruit) ash. Broad absorption at 800–1100 cm□^1^ and secondary trough at 1500–1700 cm□^1^; residual C–H features at ∼2950 cm□^1^.

**Figure 7.**
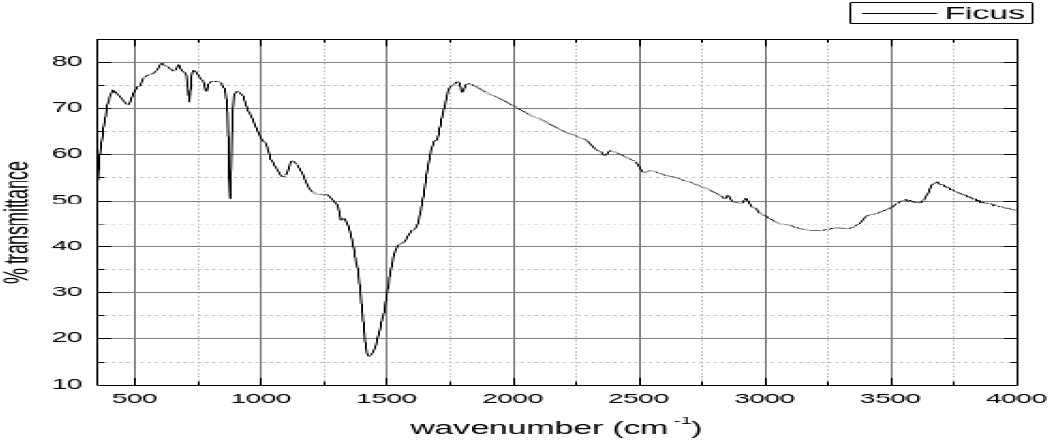
FTIR spectrum of crude aqua regia extract of Ficus benghalensis ash. The sharpest mid-IR trough (∼1400– 1500 cm□^1^; T ∼17–20%) of the vascular plant series most closely matched pure H[AuCl□].

### 3.4 UV–Visible Spectrophotometric Analysis

The UV–visible spectra confirmed the presence of [AuCl□]□ in all six species. The λmax at 372.5□nm was universally matched across all species (Table 3), representing a ∼4.5□nm bathochromic shift from the 377□nm band of pure H[AuCl□] in idealised conditions, consistent with the influence of co-dissolved organic ligands and mineral anions on the ligand-field splitting of the square-planar [AuCl□]□ complex. The 351□nm band was matched in five of six species; the 253□nm higher-energy charge-transfer transition was matched in four species.

**Table 3.** Matching UV–Visible absorption maxima (λmax, nm) between reference pure chloroauric acid and centrifuged aqua regia extracts of plant ash. +□= □present; −□=□absent.

| Ref. $\lambda_{\text{max}}$ (nm) | Acacia | Alstonia | Anacardium | Artocarpus | Ficus | Syzygium |
| --- | --- | --- | --- | --- | --- | --- |
| 377 | – | – | – | – | – | – |
| 372.5 | + | + | + | + | + | + |
| 351 | – | + | + | + | + | + |
| 324 | – | – | – | – | – | – |
| 310 | – | – | – | – | + | – |
| 308 | – | – | – | – | – | + |
| 301 | + | – | + | + | – | – |
| 294 | – | – | – | + | – | – |
| 272.5 | – | + | – | – | – | + |
| 253 | + | – | + | + | + | – |
| 221 | – | – | – | + | + | + |

### 3.5 Powder X-ray Diffraction Analysis

Powder XRD diffractograms (2θ □ 20°–80°, Cu Kα) of the heavy ash fractions of five species are presented in Figures 8–12 (duplicate *A. heterophyllus* scan excluded). All five diffractograms share a dominant, sharp reflection at ∼30° 2θ, most consistent with the calcite (104) reflection (d □ 3.035□Å, JCPDS 05-0586) or magnetite (220) at 30.1° (JCPDS 19-0629) — mineral phases expected in BIF-derived ash. The critical diagnostic cluster of reflections at 37°–50° 2θ maps directly onto the principal face-centered cubic (FCC) metallic gold reflections: Au(111) at 38.2°, Au(200) at 44.4°, and Au(220) at 64.6° (JCPDS 04-0784). This cluster was present in all five species, and its richness and intensity correlated broadly with the gold content. A prominent reflection at ∼26.6° in *A. auriculiformis* and *A. heterophyllus* is consistent with quartz (101) at 26.65° (JCPDS 46-1045), reflecting phytogenic or detrital silica in the heavy fractions.

**Figure 8.**
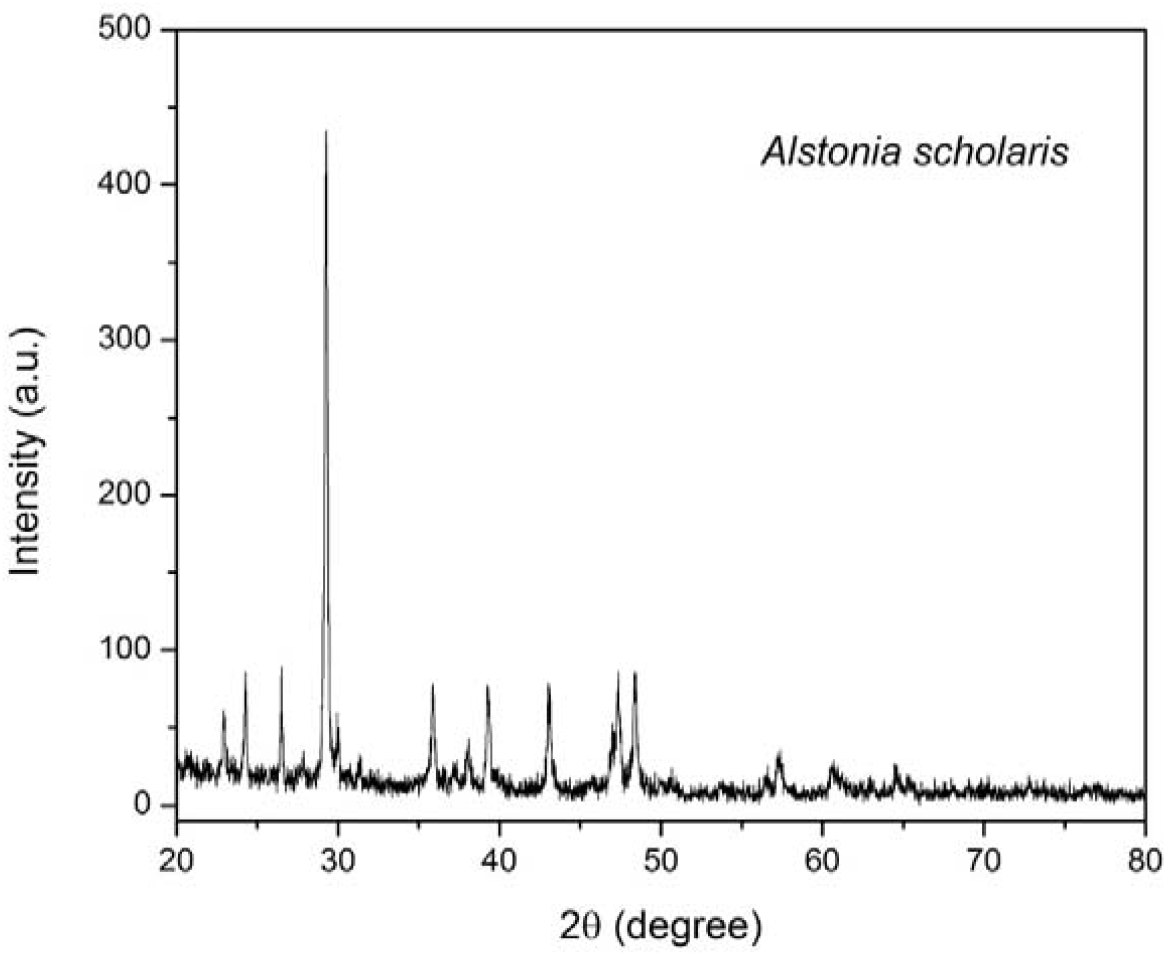
Powder XRD diffractogram of Alstonia scholaris heavy ash fraction. A dominant peak at ∼30° 2θ (calcite/magnetite) and a rich cluster at 37–50° (Au° FCC reflections). Au(111) FWHM = 0.35°; crystallite size, d = 24.8 nm.

**Figure 9.**
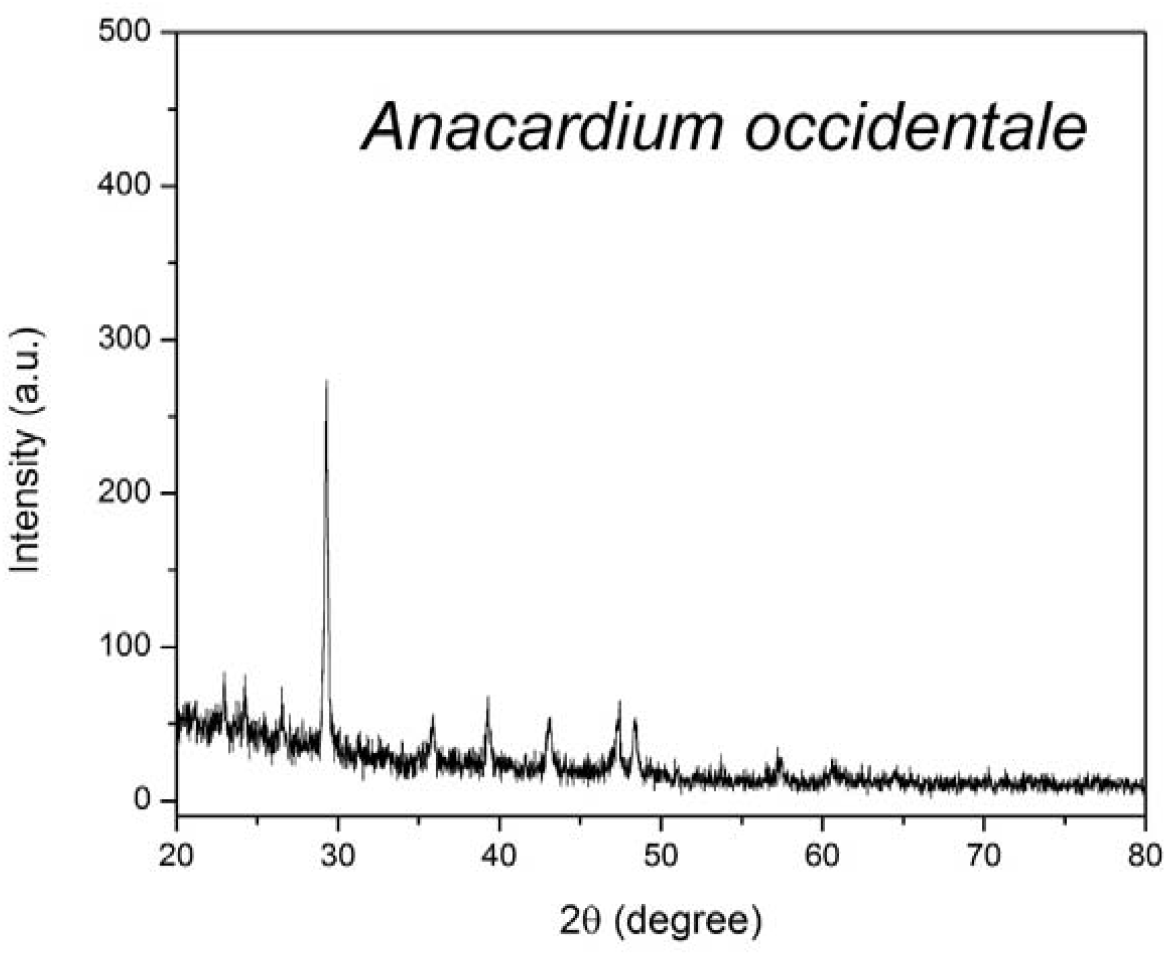
Powder XRD diffractogram of Anacardium occidentale heavy ash fraction. Elevated amorphous background at 20°–28° (biogenic opal-A/amorphous SiO□). Au(111) FWHM = 0.48°; crystallite size d = 17.7 nm (smallest value in the series).

**Figure 10.**
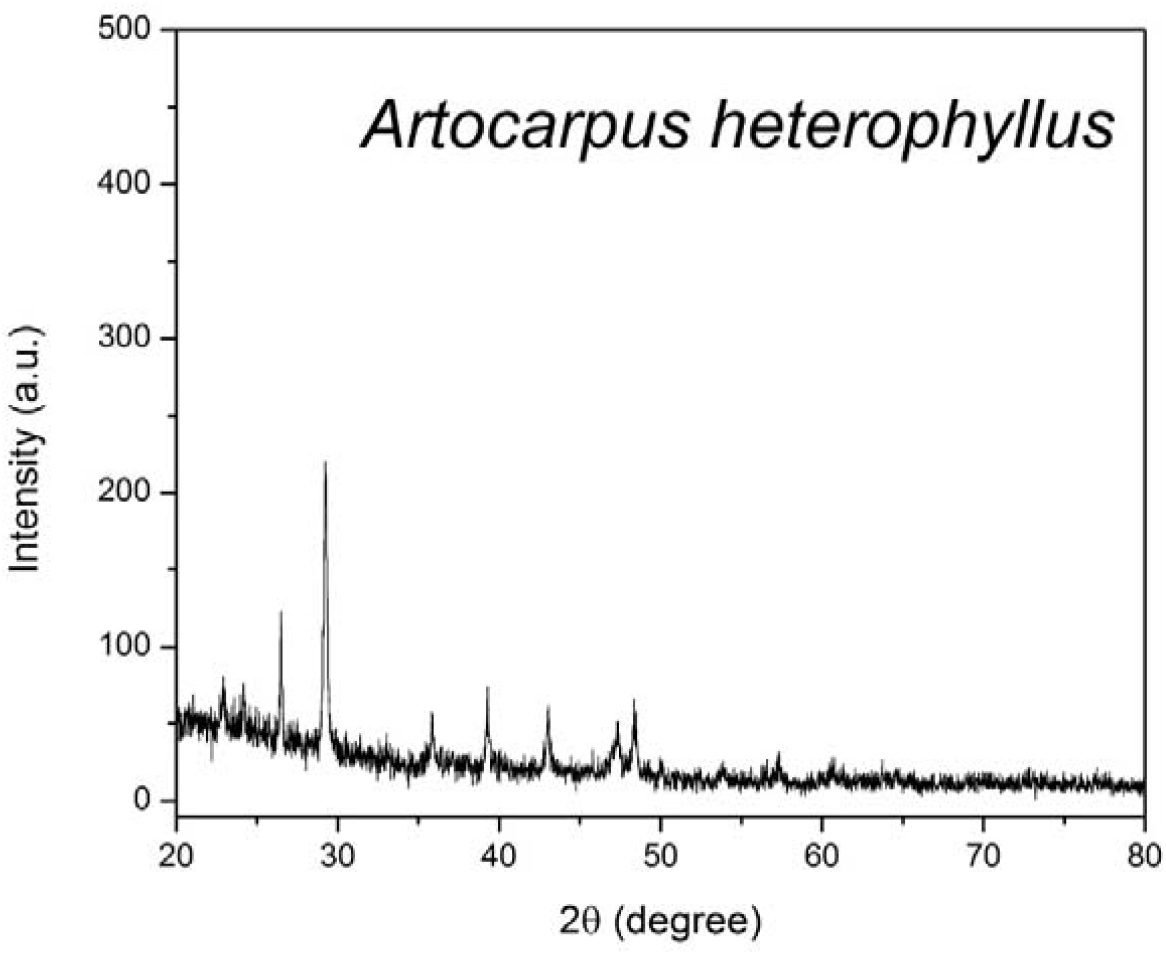
Powder XRD diffractogram of Artocarpus heterophyllus heavy ash fraction. Prominent ∼26.6° reflection consistent with quartz (101); moderate Au° cluster at 37–50°. Au(111) FWHM = 0.42°; crystallite size, d = 20.4 nm.

**Figure 11.**
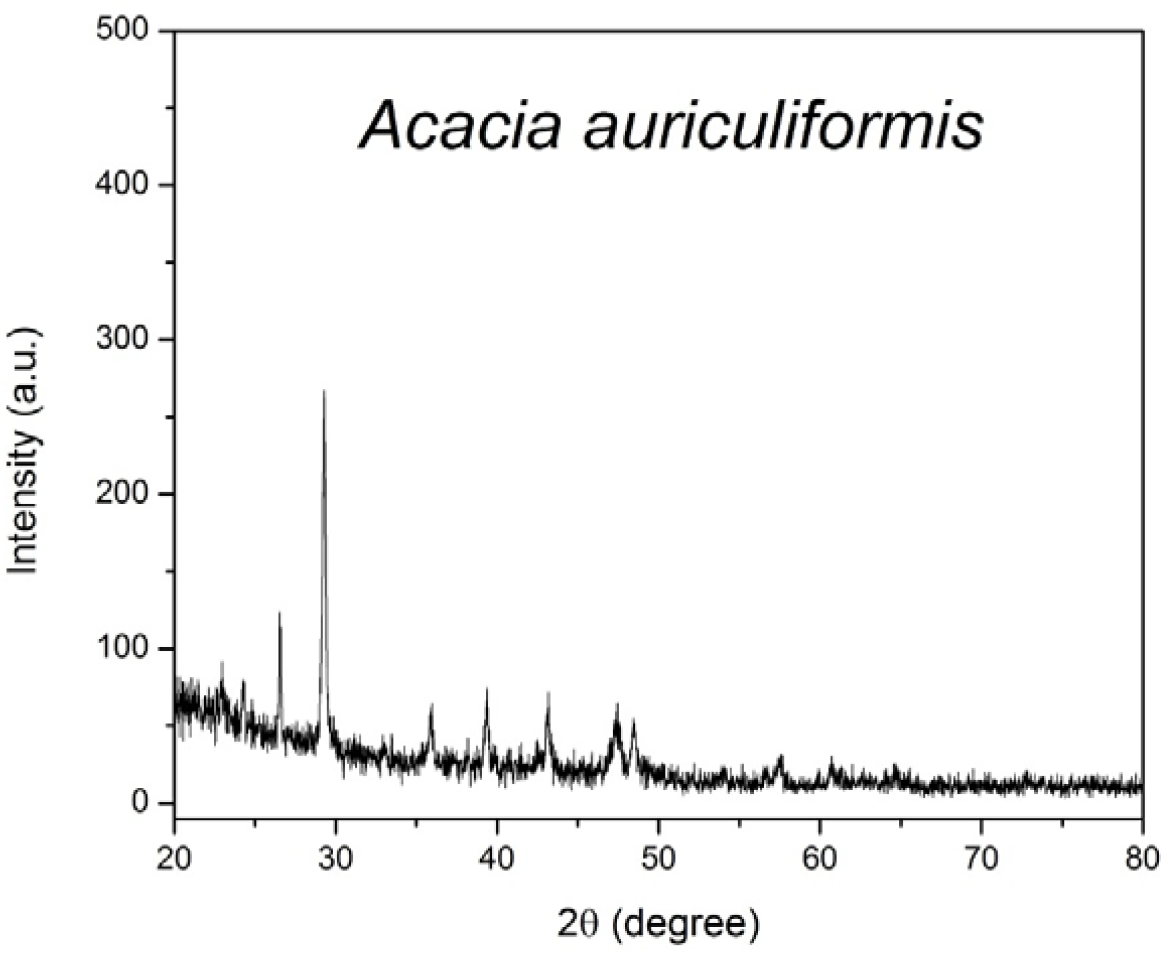
Powder XRD diffractogram of Acacia auriculiformis heavy ash fraction. A well-resolved Au° cluster at 37– 50° is consistent with the 1050 ppm gold content. Au(111) FWHM = 0.38°; crystallite size, d =; 22.7 nm.

**Figure 12.**
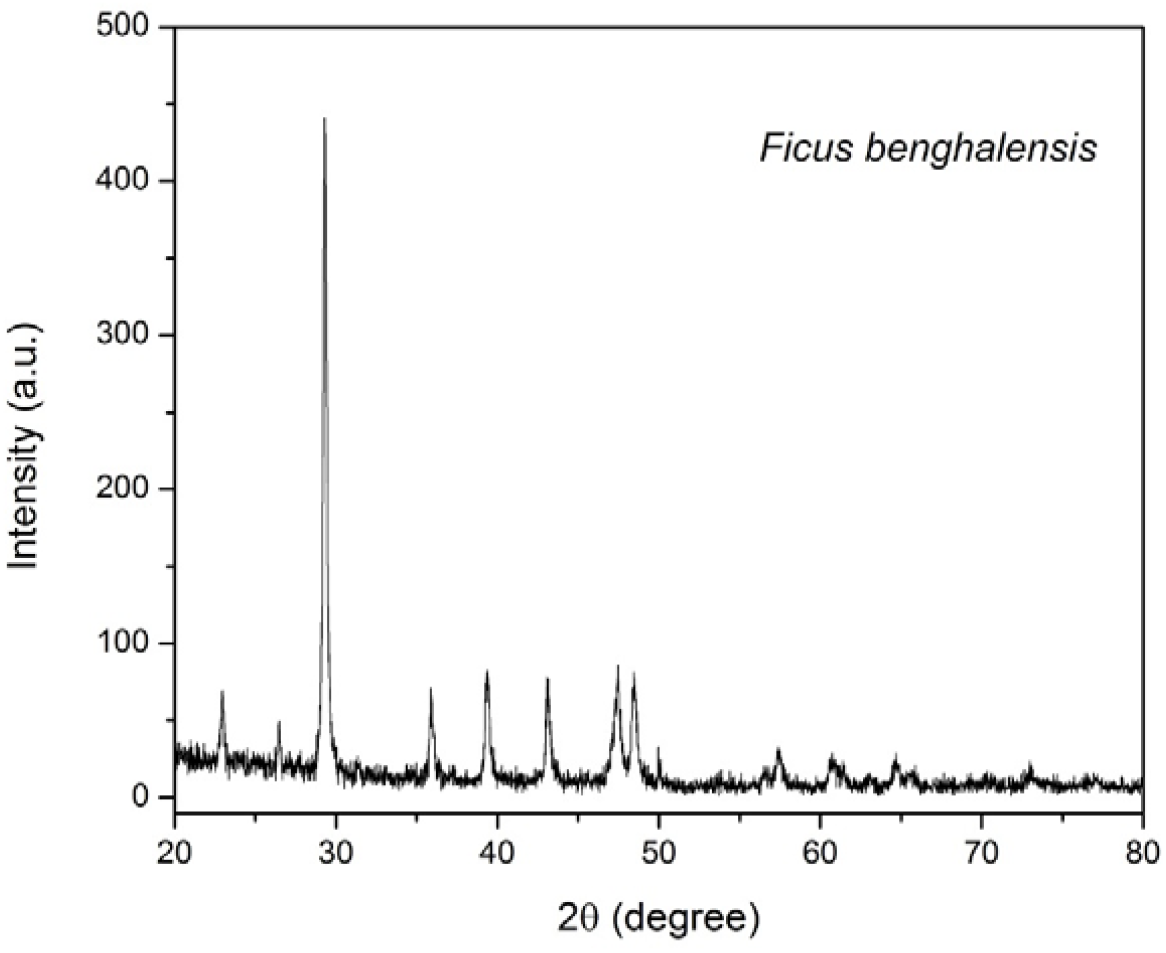
Powder XRD diffractogram of Ficus benghalensis heavy ash fraction. The sharpest and most intense dominant peak (∼30°) and the richest multiline Au° cluster (37–58°) in the series. Au(111) FWHM = 0.28°; largest crystallite size d = 31.8 nm; phytoaurolith-forming species.

### 3.6 Debye–Scherrer Crystallite Size of Phytoformic Gold

The crystallite sizes of the phytoformic Au(111) phase, calculated using the Debye–Scherrer equation from the Au(111) reflection at 2θ = 38.2°, are presented in Table 4. All values fall within the nanoparticle domain (17.7–31.8□nm), confirming that phytoformic gold in all five species consists of crystalline gold nanoparticles rather than bulk metallic gold. *Ficus benghalensis* produced the largest and most crystalline gold nanoparticles (31.8□nm), consistent with its role as the phytoaurolith-forming species. *Anacardium occidentale* yielded the smallest crystallites (17.7□nm), consistent with its poorly crystalline, amorphous-dominated ash matrix.

**Table 4.** Debye–Scherrer crystallite size of phytoformic Au(111) in heavy-ash fractions. Parameters: K □ 0.89, λ □ 1.5406 Å (Cu Kα), 2θ □ 38.2°, and βinst □ 0.10° (quadrature-subtracted).

| Species | $\beta_{\text{obs}} (^\circ)$ | $\beta_{\text{inst}} (^\circ)$ | $\beta_{\text{corr}} (^\circ)$ | $\theta (^\circ)$ | $d\text{ (nm)}$ | Au (ppm) |
| --- | --- | --- | --- | --- | --- | --- |
| *Alstonia scholaris* | 0.35 | 0.10 | 0.335 | 19.10 | **24.8** | 1100 |
| *Anacardium occidentale* | 0.48 | 0.10 | 0.469 | 19.10 | **17.7** | 675 |
| *Artocarpus heterophyllus* | 0.42 | 0.10 | 0.408 | 19.10 | **20.4** | 275 |
| *Acacia auriculiformis* | 0.38 | 0.10 | 0.367 | 19.10 | **22.7** | 1050 |
| *Ficus benghalensis* | 0.28 | 0.10 | 0.262 | 19.10 | **31.8** | 875 |

$\beta_{\text{corr}} = \sqrt{(\beta_{\text{obs}}^2 - \beta_{\text{inst}}^2)}$ ; $d$ values in bold. Sensitivity: $\pm 0.05^\circ$ in FWHM propagates to $\pm 4\text{--}8\text{ nm}$ in $d$ .

## 4. Discussion

### 4.1 Magnitude and Geological Context of Gold Accumulation

Gold concentrations of 275–1100□ppm in the heavy ash fraction are two to five orders of magnitude above crustal mean gold (∼0.004□ppm) and substantially above published phytomass concentrations in non-hyperaccumulating settings (generally <5□ppm; Girling and Peterson, 1980). This extraordinary biomagnification is attributable to the specific pedological context: the BIF/BHQ belt of North Goa constitutes an auriferous substrate, supplying anomalously high amounts of dissolved and particulate gold to the plant rhizosphere. Passive uptake via the transpiration stream, active root-mediated solubilization through organic acid exudates, and fungal-mediated mobilization in the mycorrhizal zone are all plausible contributing mechanisms (Jones et al., 1989; Christopher et al., 1999).

### 4.2 Spectroscopic Evidence: FTIR and UV–Visible

The aqua regia treatment converts elemental gold and gold-bearing mineral phases into the soluble tetrachloroaurate(III) complex H[AuCl□]. Membrane filtration (0.22□μm) removes colloidal gold and most macromolecular organic matter, yielding a filtrate dominated by ionic [AuCl□]□. The universal presence of the 1400–1700□cm□^1^ FTIR absorption envelope, the broad O–H envelope at 2800–3600□cm□^1^ characteristic of H[AuCl□]·4H□O, and the UV–Visible λmax at 372.5□nm across all six species collectively provide robust multi-technique spectroscopic confirmation of chloroauric acid formation. The degree of match with pure H[AuCl□] varied among species, with *F. benghalensis* and *A. scholaris* yielding the cleanest signatures and *A. occidentale* the most matrix-obscured spectrum, consistent with differences in ash mineral composition.

### 4.3 XRD Evidence and Nanoparticle Nature of Phytoformic Gold

The presence of Au(111), Au(200), and Au(220) FCC metallic gold reflections in all five XRD diffractograms provides direct crystallographic evidence that phytoformic gold exists as elemental, crystalline Au° in the heavy ash fraction. The Debye–Scherrer analysis definitively establishes that this gold is in the nanoparticle size range (17.7–31.8□nm), classifying phytoformic gold from these species as biogenic gold nanoparticles (bio-AuNPs) of phytogenic origin. This is a significant finding as it confirms that the biological processes of gold uptake, transport, and deposition within these species inherently produce nanoscale crystalline gold without any post-harvest chemical or physical processing.

The largest crystallites were found in *F. benghalensis* (31.8□nm), the species that also produces phytoauroliths. Larger, more ordered crystallites are consistent with the biological framework of the phytoaurolith structure, providing an organized nucleation and growth template. The smallest crystallites in *A. occidentale* (17.7□nm) and their broader, poorly resolved diffraction peaks suggest less ordered gold deposition in a matrix dominated by amorphous silica and organics from cashew shell ash. The absence of a simple correlation between crystallite size and total gold concentration (e.g., *A. heterophyllus* has 275□ppm gold but 20.4□nm crystallites, while *A. scholaris* has 1100□ppm but 24.8□nm crystallites) indicates that crystallite size reflects the biological and physicochemical conditions of gold deposition within plant tissues, not merely the total gold load.

### 4.4 Phytoauroliths: A Novel Biomineral

The designation ‘phytoauroliths’ for the strawberry-shaped isomorphic auriferous siliceous biominerals in *F. benghalensis* ash introduces a morphologically coherent new category of plant-derived gold biominerals. Phytoliths (amorphous SiO□ bodies secreted by plant cells) are well-documented in the botanical literature; phytoauroliths appear to represent a gold-enriched variant of this phenomenon, where gold deposition co-occurs with phytogenic silica secretion. Their isomorphic symmetry and consistent morphology across samples strongly suggest a biologically templated formation mechanism rather than abiotic precipitation. Full characterization by TEM-EDS, SAED, electron microprobe, and synchrotron XRF mapping is currently in progress.

### 4.5 Applied Significance

Three principal implications emerge from this study. First, all six species function as biogeochemical indicator species for gold anomalies in the BIF terrain, enabling low-cost, non-invasive prospecting through systematic ash analysis of litter. Second, the confirmed nanoparticle nature of phytoformic gold opens a direct route to bio-AuNP production: ashing followed by controlled extraction could yield size-selected gold nanoparticles of phytogenic origin without the chemical reducing agents, stabilizers, or energy-intensive synthesis conditions required by conventional wet-chemistry nanoparticle protocols. Third, the species demonstrate phytoextraction potential for gold recovery from low-grade ore or mining waste, with *A. scholaris* and *A. auriculiformis* as the highest-accumulating candidates (1100 and 1050□ppm respectively).

## 5. Conclusions

This study established the following:

1. The gold content in the heavy ash fraction ranged from 275 to 1100 ppm across the six species, with a biomagnification factor of ∼70,000–275,000 over the crustal background.
2. FTIR spectroscopy confirmed [AuCl□]□ as the dominant dissolved gold species in 0.22 μm filtered aqua regia extracts of all six species.
3. UV–visible spectroscopy independently confirmed the formation of chloroauric acid with λmax = 372.5 nm across all six species.
4. Powder XRD identified FCC metallic Au° reflections [Au(111), Au(200), Au(220)] in all five species analysed, confirming crystalline elemental gold in the heavy ash fraction.
5. Debye–Scherrer analysis of the Au(111) reflection established crystallite sizes of 17.7–31.8 nm, confirming that phytoformic gold exists as biogenic gold nanoparticles (bio-AuNPs) in all the species.
6. The novel biomineral category ‘phytoauroliths’ — strawberry-shaped isomorphic auriferous siliceous biominerals — was discovered in *Ficus benghalensis* ash, corresponding to the largest crystallite size (31.8 nm) in the series.
7. The described protocol — ash, aqua regia extraction, membrane filtration, and UV– Vis/FTIR/XRD confirmation — constitutes a validated low-cost method for biogeochemical gold anomaly detection and forms the basis of a patentable prospecting and bio-AuNP production system.

## Acknowledgements

The authors thank the UGC-SAP Phase-II Program on Biodiversity, Bioprospecting, and Biotechnology, and the RNSB Group for financial and infrastructural support. Goa University is acknowledged for providing laboratory facilities.

## Author Contributions

**Naik Shruti**: Sample collection, ash preparation, spectroscopic and XRD data acquisition, and data analysis. **Kamat N.M**.: Concept, study design, Debye–Scherrer analysis, spectral interpretation, manuscript drafting and revision, project supervision. Both authors have read and approved the final manuscript.

## Data Availability

Raw spectral data (FTIR, UV–Vis), XRD diffractograms, all figures, and the manuscript are permanently archived on Zenodo: https://doi.org/10.5281/zenodo.20213980 (Naik & Kamat, 2026, v1.0, CC BY 4.0). All data supporting the conclusions are presented in this manuscript.

## References

Christopher, L., Baker, W.J., and Gemmell, J.B. (1999). Gold uptake by plants. Gold Bulletin, 32(2), 48–52. 10.1007/BF03214784

Dessai, A.G. (1999). Geology and mineral resources of Goa. Geological Survey of India, Miscellaneous Publication, 30 (Part IV-4), 1–45.

Girling, C. A., & Peterson, P. J. (1980). Gold in plants. Gold Bulletin, 13(4), 151–157. 10.1007/BF03216550

Jones, H.C., Blaylock, M.J., and Chaney, R.L. (1989). Gold uptake by perennial ryegrass: the influence of humates on gold cycling in soil. Biogeochemistry, 7(1), 3–10. 10.1007/BF00000885

Naik, S. and Kamat, N.M. (2026). Phytoformic Gold in Ash Samples of Plants from the North Goa Iron Ore Mining Belt: Detection, Characterisation, X-ray Diffraction, and Spectroscopic Evidence for Biogeochemical Gold Nanoparticle Formation (v1.0). Zenodo. 10.5281/zenodo.20213980

Scherrer, P. (1918). Bestimmung der Größe und der inneren Struktur von Kolloidteilchen mittels Röntgenstrahlen. Nachrichten von der Gesellschaft der Wissenschaften zu Göttingen, 26, 98–100.

